# New soft tissue data of pterosaur tail vane reveals sophisticated, dynamic tensioning usage and expands its evolutionary origins

**DOI:** 10.1101/2024.07.01.601487

**Authors:** Natalia Jagielska, Thomas G. Kaye, Michael B. Habib, Tatsuya Hirasawa, Michael Pittman

## Abstract

Pterosaurs were the first vertebrates to achieve powered flight. Early pterosaurs had long stiff tails with a mobile base that could shift their center of mass, potentially benefiting flight control. These tails ended in a tall, thin soft tissue vane that would compromise aerodynamic control and efficiency if it fluttered excessively during flight. Maintaining stiffness in the vane would have been crucial in early pterosaur flight, but how this was achieved has been unclear, especially since vanes were lost in later pterosaurs and are absent in birds and bats. Here we use Laser-Stimulated Fluorescence imaging to reveal a cross-linking lattice within the tail vanes of early pterosaurs. The lattice supported a sophisticated dynamic tensioning system used to maintain vane stiffness, allowing the whole tail to augment flight control and the vane to function as a display structure.

## Introduction

Pterosaurs were the first vertebrates to achieve powered flight (Palmer 2017). The first pterosaurs, the non-pterodactyloids, had long, stiff tails with a mobile base (Frey, Tischlinger et al. 2003), similar to some dinosaurs like *Velociraptor* (Persons IV and Currie 2012). Many of these tails end in a soft tissue ‘vane’ (Marsh 1882, Döderlein 1929, Frey, Tischlinger et al. 2003) (Fig. 1), which may have contributed to passive stability in flight. A primary role in display has also been suggested (O’Brien, Allen et al. 2018), given ontogenetic shape changes in the vane and the fact that, unlike most aircraft, flying animals do not need vertical control surfaces to be yaw-stable during turns (Bowers, Murillo et al. 2016). The vanes have been interpreted as steering aids (Frey, Tischlinger et al. 2003). The length and stiffening of the tails suggest that they might have been important in early pterosaurs for control based on mass shifting or inertial control, as purported for terrestrial theropods with convergent tails (Persons IV and Currie 2012). Such dynamic control could greatly improve maneuverability and/or stability. However, vane fluttering would be extremely costly and destabilising unless the vane was tensioned while under aerodynamic load. Tail vanes feature thick, evenly spaced, internal structures roughly perpendicular to the caudal series (Döderlein 1929), that are said to resemble neural spines and haemal arches (Marsh 1882). These structures are presumed to have minimised fluttering and prevented buckling in the same way that spars, ribs, stringers, and longerons do in airplane wings and tail-fins, but others have proposed that they were flexible and cartilaginous (Marsh 1882), especially since their preserved appearance varies. Here we use Laser-Simulated Fluorescence (LSF) imaging of *Rhamphorhynchus* specimens from the Upper Jurassic Solnhofen Limestones (Kaye, Falk et al. 2015, Pittman, Barlow et al. 2021) to investigate the vane’s structural properties, explore its usage, its evolutionary origins and the context for its disappearance in later pterodactyloids (Frey, Tischlinger et al. 2003).

**Figure 1.**
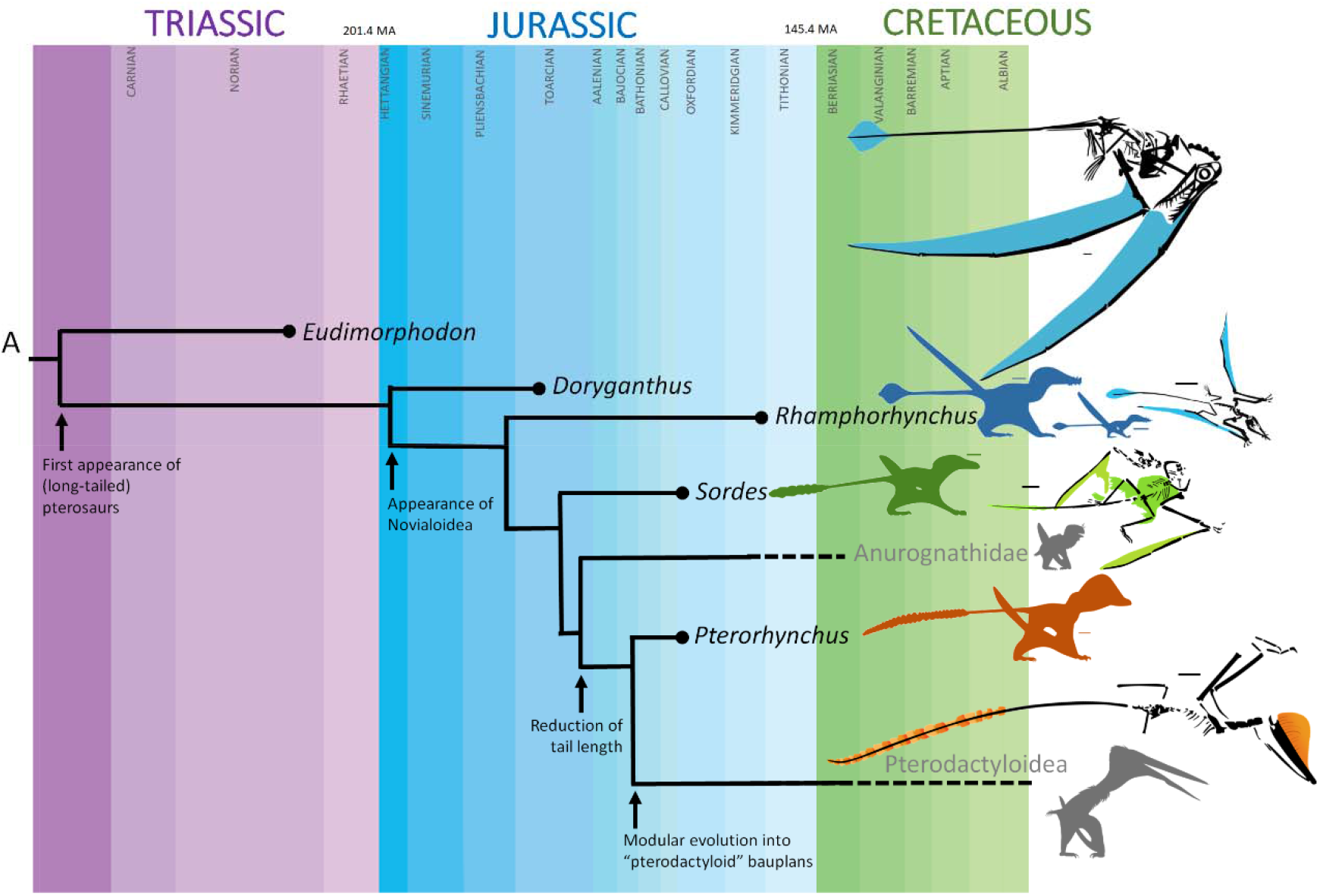
Long-tailed early-diverging non-pterodactyloid pterosaurs had diverse tail vanes but these disappeared in later-diverging short-tailed pterodactyloids. Blue, ontogenetic morphs of *Rhamphorhynchus muensteri*: BSPG 1938 I 503a and ROM VP 55352. Green, *Sordes pilosus* PIN 2585/3. Orange, *Pterorhynchus wellnhoferi* CAGS02-IG-guasa-2/DM608. Scale bars are 3 cm.

## Results

Over 100 Solnhofen pterosaur fossils were examined for well-preserved tail vanes using an ultraviolet torch. Four exceptional specimens were then imaged under Laser-Stimulated Fluorescence (LSF). Three specimens exhibited tail vanes under white light but the vane of NHMUK PV OR 37003 was only visible under LSF. LSF confirmed the soft tissue extent of the vanes and revealed hidden anatomical details, especially in NHMUK PV OR 37003 and 37787 and NMS G.1994.13.1 (Fig. 2A-C), where vane areas fluoresced pink and white, indicating soft tissue preservation (Pittman, Barlow et al. 2021). Tail vanes are sub-symmetrical and diamond-shaped in NHMUK PV OR 37003 and 37787 and NMS G.1994.13.1 with a length of 700 mm, 750 mm and 720mm making up 21%, 22% and 21% of the total tail length of 320 mm, 362 mm and 348 mm respectively (Fig. 2). At its widest point, about two-thirds along its length, the vane is 41 mm across in NHMUK PV OR 37003 and 37787 but the widest in NMS G.1994.13.1 (55 mm), even wider than in BSP 1907 I 37 (46 mm) and almost twice as wide as YPM VP 1778 (30 mm). Under LSF, partial edges of the vane are visible, along with at least 17 relatively straight structures in NHMUK PV OR 37787 (10+ in NHMUK PV OR 37003 and 11+ in NMS G.1994.13.1); projecting vertically, near-perpendicularly to the tail skeleton (based on position of chevron bones e.g., NHM PV OR 37787). In NHMUK PV OR 37003 and 37787 and NMS G.1994.13.1, these are relatively thick (0.6 - 1 mm) and appear to be hollow, suggesting they were rod-like, and were arranged in parallel ∼3 - 8 mm apart. In NHMUK PV OR 37003 and 37787 they are rarely preserved dead- straight, but are straighter in NMS G.1994.13.1, especially anteriorly. In YPM VP 1778 and BSP 1907 I 37 the vertical structures show more pronounced undulations giving them a sigmoidal morphology. To our knowledge, in NHMUK PV OR 37003 alone, there is a second layer of thinner and more numerous fibres that run across the thick vertical structures, subparallel to the long axis of the tail and become more and more closely spaced as they reach the tail tip. Together, the vertical structures and subhorizontal fibres form a cross-linked lattice. The thick outer margin of the vane is undulated in dorsal view in both NHMUK PV OR 37003 and 37787, with a trough ∼0.7 mm deep where the thick vertical structures meet the margin and a convex peak roughly mid-way between each pair of vertical structures. A similarly thick outer margin is also observed anteriorly on the tail vane of NMS G.1994.13.1 and BSP 1907 I 37.

**Figure 2.**
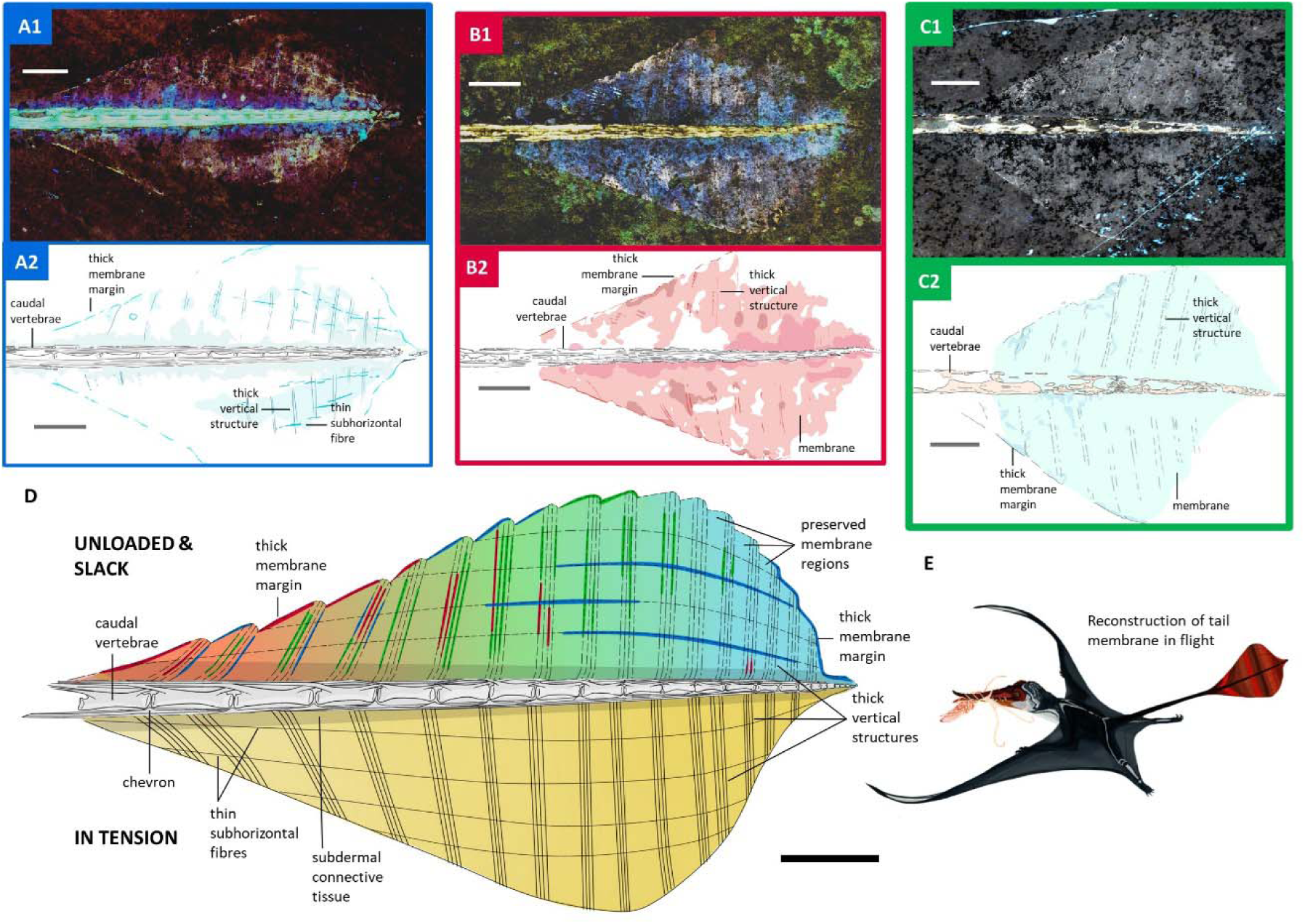
Tail vane of *Rhamphorhynchus muensteri*. **A1**. LSF image of NHMUK PV OR 37787. **A2**. Line drawing of LSF image of NHMUK PV OR 37787. **B1**. LSF image of NHMUK PV OR 37003. **B2**. Line drawing of LSF image of NHMUK PV OR 37003. **C1**. LSF image of NMS G.1994.13.1. **C2**. Line drawing of LSF image of NMS G.1994.13.1. **D**. Interpretative line drawing of *Rhamphorhynchus muensteri* tail vane unloaded and slack as well as in tension. Combines LSF results of NHMUK PV OR 37003 and 37787 as well as NMS G.1994.13.1 (A1-C2). **E**. Life reconstruction of *Rhamphorhynchus muensteri* using its tail vane during flight. All scale bars are 1 cm.

## Discussion

In order to discuss the tail vane’s structural properties and their implications, we will first introduce the relevant biomechanical concepts and how they have previously been applied to pterosaur wings. Material stiffness is a measure of a material’s resistance to deformation and can be summarised by a variable known as the modulus of elasticity (E) which is the ratio of stress (σ) to strain (L) during deformation. Stress, is the applied force divided by the area over which it is exerted, while strain refers to the change in length of the material: positive strain is stretching (tensile strain) and negative strain is compression (compressive strain). E can differ substantially for different kinds of loading - in soft tissues, resistance to tension is usually much greater than resistance to compression. For instance, “true” ligaments running between bones typically only load in tension (i.e., when stretched), they essentially just “fold up” if compressed (Woo, Nguyen et al. 2007). Stiffer materials have a higher value of E than flexible materials: it takes more stress to produce the same strain (they deform less for a given stress).

When the soft tissue surface of the pterosaur wing was placed under aerodynamic load, the associated stress was expected to have stretched the wing surface so it curved towards the side of lower air pressure. This resulted in “auto-cambering” of the wing surface, which became more concave beneath (and more convex above). The degree of auto-cambering was limited by stiffening mechanisms preventing excessive stretching (Palmer and Dyke 2012). Stiffening was provided by a combination of active mechanisms through muscles within the wings as well as passive mechanisms, including keratinisation (stiff keratins have the highest E of any soft tissue) and reinforcement by fibres within the wing (actinofibrils) (Palmer and Dyke 2012). Actinofibrils reinforced the wing against stretching by having a comparatively high value of E in tension: as the entire wing surface bowed, both the ventral and dorsal surfaces were loaded in tension, which applied a tensile load to the actinofibrils. The actinofibrils stretched according to their elastic modulus in tension, which limited the stretching of the entire wing surface. This stands in contrast to the cantilever bending of a hard tissue “beam” such as a long bone, where one side is loaded in compression and the other side is loaded in tension.

The pterosaur tail vane would have experienced asymmetric air flow anytime it was deflected (even slightly) to the right or left during flight. This would produce a moderate amount of lift, oriented laterally, to either the right or left side depending on the motion of the tail. While this modest lift would have been of little use for flight control, it would have had a tendency to stretch the vane to one side or the other out of plane, much like the auto-cambering of the pterosaur wing (Palmer and Dyke 2012). The associated, rapid, stretching and release this would cause is known as aeroelastic flutter (Lillico, Butler et al. 1997), and it produces extremely high drag coefficients. Drag on the tail vane would therefore be substantially reduced if the tail vane had a mechanism of tensioning.

Our investigation of the tail vane under LSF revealed a lattice of two sets of structures: one set are thicker, regularly-spaced tube-like structures with a vertical long axis (Fig. 2). The second set are thinner, more numerous subhorizontal fibres that criss-cross the thick vertical structures (Fig. 2). We propose that they both worked as tensile-loaded structures similar to actinofibrils in the wings, but acting in tandem due to the lattice cross-links to resist stretching in multiple directions. This is analogous to how a fabric with fibres running in multiple directions resists stretching. The tail vane would have stretched in the direction of mediolateral deviation. Since the surface would be stretched, the distances between the lattice fibres would increase - pulling, especially, the thicker vertical structures apart. However, these movements would have been limited by both the bending strength of these relatively large vertical “tubes” and, more importantly, the cross-linking subhorizontal fibres. Since the subhorizontal fibres linked across the vertical structures, any stretching of the vane that spread the vertical structures apart from each other would stretch the subhorizontal fibres along their long axis – loading them in tension. This load would ultimately be taken up at the tail tip, where the fibres converged. This loading regime is consistent with the concavities along the vane edge being aligned with each structure tip, indicating that, under load, the spaces between the structures stretched until the outer edge was linear (Fig. 2D), with the fossilised position preserving the unloaded slack condition (Fig. 2D). We see no evidence of active tensioning capacity using muscle and therefore posit that this soft tissue lattice was the primary stiffening adaptation for the tail vane. We are unaware of any extant organisms with an analogous functional feature.

As the precise material composition of the preserved lattice remains elusive, we do not know the absolute value of E for the observed structures, and therefore, cannot calculate expected absolute deflections of the tail vane. However, given the orientation and apparent density of the structures, we can confidently predict that the presence of the lattice would increase the overall elastic modulus of the tail vane in tension, probably substantially. For perspective, even if all of the observed soft tissue was collagen in life, thicker, more condensed collagen fibres can have a tensile E 3.6 times that of thinner, less compact fibres (Wenger, Bozec et al. 2007). To be visibly distinct under LSF, the density and thickness of the structures in the lattice must have been several times that of the surrounding soft tissue matrix, so it is reasonable to surmise that the lattice could have stiffened the vane by a large margin - probably an order of magnitude, at least.

Thus, our results suggest that the tail vane maintained effective stiffness dynamically with internal tension of a cross-linked lattice that minimised excessive vane flutter and associated drag production. This structural integrity would have permitted the vane to be an effective display structure without incurring excessive aerodynamic costs (Fig. 2E). This tensioning would also have allowed the tail, as a whole, to be used for mass-shifting-based aerodynamic control without incurring adverse effects of a fluttering vane during rapid tail motions. Tail vane shape changes through ontogeny (Bennett 1995, O’Brien, Allen et al. 2018) and between species (Marsh 1882, Döderlein 1929, Frey, Tischlinger et al. 2003), which underscores the importance of the tail vane in early pterosaur evolution. As pterodactyloid pterosaurs evolved a shorter body plan with an anterior center of mass and large heads with cranial crests as the primary display structures (Jagielska and Brusatte 2021), both the control and display functions of the tail were absorbed by the wings and head.

The new soft tissue information also provides clues about the evolutionary origins of the tail vane itself. The cross-linked lattice recognised in this study suggests that the tail vane of early pterosaurs developed from a single contiguous structure rather than a combined structure of scales or feather-like integuments. While the undulating vane edge (Fig. 2D) might reflect an epidermal patterning, the internal part of the tail vane was likely filled with connective tissue underneath the epidermal layer. The medial part of the tail vane, namely the periphery of the caudal vertebrae, has a different tone under fluorescence and the vertical structures lose clarity when compared to the lateral part of the tail vane (Fig. 2A, B). This potentially indicates a thicker subdermal connective tissue surrounding the caudal vertebrae. Therefore, the tail vane of pterosaurs consisted of bilateral fleshy folds on the end of the tail, comparable to the cetacean fluke that envelopes dense connective tissue (Gavazzi, Nair et al. 2024). The growth series of the tail vane shape in *Rhamphorhynchus muensteri* begins with an extended teardrop/oval shape, becoming diamond-shaped (Fig. 2A, B), and eventually triangular (Bennett 1995). These three shapes parallel the shape changes of the cetacean fluke during embryonic development (Štěrba, Klima et al. 2000). It is possible that both the pterosaur tail vane and the cetacean fluke evolved through a shared developmental mechanism, perhaps a co-option of the signaling pathway that drives appendage outgrowth (Gavazzi, Nair et al. 2024), eventually bringing about improved fluid dynamics of the limbs.

## Materials and Methods

Over 100 Solnhofen pterosaur fossils were examined for well-preserved tail vanes using an ultraviolet torch at the Bayerische Staatssamlung für Paläontologie (Munich), Museum für Naturkunde (Berlin), Jura Museum (Eichstätt), Natural History Museum (London), National Museum of Scotland (Edinburgh) and Royal Ontario Museum (Toronto) representing the pterosaur inventories in their care. These specimens were screened without bias regardless of their ontogenetic stage and preservation (i.e., completeness, degree of articulation and the presence or absence of soft tissues under white light). Out of this significant sample, only four specimens were found to preserve tail vane soft tissues, which were then imaged under Laser-Stimulated Fluorescence (LSF) (Kaye, Falk et al. 2015) at the Natural History Museum, National Museum of Scotland, and the Royal Ontario Museum. LSF involved projecting a 405 nm violet laser diode from a line lens and scanning it over the specimens in a dark room following standard laser safety protocol. Long exposure photographs over 30 seconds were taken with a Nikon digital single-lens reflex camera fitted with a 425 nm laser blocking filter. LSF images were then postprocessed for equalisation, saturation, and colour balance across the entire images in Adobe Photoshop CS6. For more details see Extended Methods in Supplementary File 1.

## Data Availability

All relevant data is provided in the manuscript and Supplementary File 1.

## Acknowledgments

Mike Day, Nick Fraser, David Evans and Kevin Seymour are thanked for granting study access to specimens in their care.

## Author Contributions

Conceptualisation, M.P., N.J. and T.G.K; Methodology, M.P., T.G.K., M.B.H. & N.J.; Validation, M.P., N.J., T.G.K. & M.B.H.; Formal Analysis, M.P., N.J., M.B.H., T.H. & T.G.K.; Investigation, N.J., M.P., T.G.K., T.H. & M.B.H.; Resources, M.P. and T.G.K.; Writing – Original Draft, M.P., N.J., M.B.H. & T.G.K.; Writing – Review and Editing, M.P., N.J., M.B.H., T.H. & T.G.K.; Supervision, M.P.; Project Administration, M.P.; Funding Acquisition, M.P.

