## Supplementary File 1 for "New soft tissue data of pterosaur tail vane reveals sophisticated, dynamic tensioning usage and expands its evolutionary origins"

**Extended Methods**

*Rhamphorhynchus muensteri* NHMUK PV OR 37003

Initially described by von Meyer (1846) as *R. gemmingi*; followed by *R. meyeri* (Owen, 1870), described under Nr. 39 in Wellnhofer, 1975; synonymised under the same species by Bennett, 1995. No keratin appears conserved on pedal digits or rhamphotheca, despite good preservation. Wellnhofer in his assessment does not mention the presence of a tail vane and the feature is not discernible in visible light. Detected with UV and then imaged more vividly under LSF. The specimen is better articulated than NHMUK PV OR 37787. Ontogenetically, on basis of the size of preserved elements and tail vane morphology, it likely belongs to a similar growth stage to NHMUK PV OR 37787. The distal end of tibia is unfused, pointing at immaturity of the animal.

The slab contains a ventrally exposed lower jaw and heavily disarticulated postcranial material. It preserves fragments consisting of disarticulated flight apparatus and partially preserved scapula-coracoid. 11 rib-bearing dorsal vertebrae in articulation; disarticulated unfused pubic elements, both hindlimb elements with paired well-preserved tarsals in articulation. Metacarpal IV is 23 mm long. The wing phalanges make up an incomplete ~380 mm long wing (WP1 = ~113 mm but incomplete, WP2 = ~112 mm but incomplete, WP3 = ~101 mm, WP4 = ~54 mm but incomplete). The tibia is 52 mm long and the incomplete femur ~32 mm long. Metatarsals 1-5 are 26 mm, 28 mm, 27 mm, 21 mm and 10 mm long respectively.

The preserved tail vane is slightly shorter than lower jaw (89 mm but incomplete), with two features being similar sized. The total tail length is 320 mm, it is preserved with filiform and almost entirely intact, only disarticulating anteriorly. The terminal tail vane is 70 mm of the tail length (22%). The vane encompasses over fifteen vertebrae and is 41 mm wide.

*Rhamphorhynchus muensteri* NHMUK PV OR 37787

Assigned *Rhamphorhynchus muensteri* by Goldfuss, 1831; followed *by R. gemmingi* by von Meyer, 1846; the specimen was then re-assessed under label Nr. 42, in Wellnhofer, 1975; and cumulatively assigned *R. muensteri* as all *Rhamphorhynchus* species were synonymised as different year classes of same species (Bennett, 1995). Wellnhofer already noted the faint preservation of a tail vane in his assessment.

The slab contains largely disarticulated postcranial material. The skull is partially preserved but severely deformed and eroded, with an outline of the paired ramus (70 mm rami length) and upper crania, retaining anterior dentition with larger teeth (14 mm) displaced anteriorly to the preserved jaws. Most postcranial material is composed of the disarticulated ribcage, with scattered ribs and a partially articulated dorsal vertebral segment. Element of the very deformed sternum and paired scapula-coracoids are visible along with a humerus. The humerus is 39 mm long, a stocky bone with deflecting deltopectoral crest characteristic of *R. muensteri*. The radius is 71 mm long and fourth metacarpal is 24 mm long. The wing phalanges make up a ~465 mm long wing (WP1 = 123 mm, WP2 = ~119 mm, WP3 = ~112 mm, WP4 = 111 mm), accumulating to a wingspan of ~1.2 metres, with the terminal phalange retaining a 165° curvature.

The vertebral column is preserved as modular chunks in partial articulation, composed of presacral vertebrae, sacral vertebrae and over 30 caudal vertebrae in partial articulation; the filiform of the caudal section disarticulate and splay anteriorly. Orientation of preservation enables good visibility of individual caudal bones, as in some specimens these are obscured by elongate, onlapping chevrons and zygapophyses. The total tail length is 362 mm. The vane makes up 75 mm of the tail (21%) and is 41 mm at its widest point.

*Rhamphorhynchus muensteri* NMS G.1994.13.1

The specimen was assessed by Wellnhofer (pers. comm.) and mentioned in Frey *et al.* (2003) for the likelihood of preserving a “throat pouch”. The specimen was later used in “tail fin” calculations by O’Brien *et al*., 2018, but remained formally undescribed. While retaining a well-preserved tail vane, the wing membrane, despite discolouration and topographic lineation, does not preserve actinofibrils. Keratinous terminations of rhamphotheca and unguals are also absent. The skull has a proportionally sizeable orbit while the wing phalange extensor process and pubic fusing point to a late-stage sub-adult ontogenetic stage.

The slab contains a largely complete and modularly articulated cranial and postcranial skeleton. The skeleton is preserved contorted, seemingly “biting its tail” with cervical and dorsal vertebrae falling severely out of articulation. Its well-preserved skull, including the sclerotic ring, is 112 mm long with a 86 mm long lower jaw. The scapula-coracoid pair are preserved separately, with forelimb elements (humerus, radius, ulna, metacarpal, manual phalanges) severely deformed or missing. The sterna is ~30 mm. Wings preserve all phalanges, measuring ~467 mm in total length (WP1 = ~130 mm; WP2 = ~134 mm; WP3 = ~125 mm; WP4 = ~78 mm). Its femur and metatarsals are obscured, with only 64 mm measurable on the tibia.

While experiencing some deformation, the tail stems from the terminal sacrum and is fully preserved in articulation. The tail measures 348 mm, out of which ~72 mm is taken up by the tail vane. Making up 21% of the total tail length. The tail vane is “stockier” in comparison to the NHMUK specimens, having a wider anterior opening and measuring 55 mm at its widest section. The vane encompasses at least 15 caudal vertebrae.

**Additional Comparative Anatomy**

The vanes of NHMUK PV OR 37003 and 37787, NMS G.1994.13.1 and YPM VP 1778 form anterior, lateral and posterior angles of 40°, 45° and 90°; 107°, 113° and 105°; 104°, 100° and 90°; 44°, 107° and 80°, respectively. NHMUK PV OR 37003 and 37787 are more faintly distinguishable in visible light by localised discolouration compared to NMS G.1994.13.1 and YPM VP 1778. NHMUK PV OR 37003 and 37787 and NMS G.1994.13.1 have tail vanes occupying a similar proportion of the tail (23% and 22%) as BSP 1907 I 37 (74 mm) and YPM VP 1778 (50 mm). The vertical tail vane structures make an angle of ~90° with the tail axis at the widest point of vane with a more acute angle of 57°, 65°, 66° and 57° in NHMUK PV OR 37003 and 37787, NMS G.1994.13.1 and YPM VP 1778 respectively. There is no marked change in caudal morphology within and outside the periphery of the tail vane. The vane encompasses at least 15 caudals in NHMUK PV OR 37003 and NMS G.1994.13.1 and at least 17 in NHMUKPOV OR 37787, out of which, 13 appear to bear elongated zygapophyses in 37703 and at least 14 in 37787 (16 caudals within YPM VP 1778 and BSP 1907 I 37). The vane is extrapolated to originate from the anterior end of a caudal in NHMUK PV OR 37003 and 37787 and NMS G.1994.13.1. In NHMUK PV OR 37003 and OR 37787 and NMS G.1994.13.1, these thick vertical structures stem from caudal articulation points in the distal portion of the vane but anteriorly there is no obvious pattern. These data do not support their association with the neural spines and haemal arches (*contra* Marsh (1882)).
